# Personality homophily drives female friendships in a feral ungulate

**DOI:** 10.1101/2024.06.23.600227

**Authors:** Debottam Bhattacharjee, Kate J. Flay, Alan G. McElligott

## Abstract

Friendships, exhibited by both humans and non-human animals, have considerable adaptive benefits. In humans, similarity or homophily in personality is considered a proximate mechanism driving friendships, yet little is known about the behavioural ‘decision rules’ underlying animal friendships. Some empirical research suggests that animal friendships can be driven by personality homophily. However, these studies are restricted to non-human primates, limiting our understanding of the mechanisms of friendships. We investigated a feral population of water buffalo (*Bubalus bubalis*) to determine whether homophily in personality drives friendships in this ‘non-model’ social species in free-ranging environmental conditions. We conducted observations on females (*n=30*) and assessed their friendships and personalities. Close spatial proximity served as a behavioural indicator of friendship, validated by affiliative body contact. An objective ‘bottom-up’ method revealed three personality traits – *social tension*, *vigilance*, and *general dominance*. We found that females with comparatively lower personality differences in social tension and general dominance traits exhibited significantly higher close spatial associations. We did not find an effect of kinship on close spatial associations. Our findings show that friendships in buffalo can form based on personality homophily, a decision rule attributed predominantly to primates. We discuss these findings in light of buffalo socioecology but emphasise their implications in the broader evolutionary context of animal personalities and friendships.

## Background

In social species, including humans, the strength of affiliative ‘ties’ in a social network is not homogenous, indicating varying degrees of relationships among individuals. Preferential strong social associations or friendships are of particular interest as they positively correlate with health, well-being, and survival benefits [1, 2]. Although conceptualised relatively recently in non-human animals (hereafter, animals) [3], growing convergent evidence suggests that human-like friendships can form in animals [4, 5]. Similar to humans, animal friendships can be stable, long-lasting [6, 7], and form beyond the extent of kin relationships [8]. The primary evolutionary explanation of friendship lies in their low levels of uncertainty, where, in comparison to a non-friend, a friend provides assured fitness benefits, and these benefits surpass the costs of maintaining friendships [9]. For instance, friendships foster cooperation, which minimises the probability of ‘cheating’ [10–13]; the strong emotional underpinning of friendship can act as a ‘social buffer’ during aversive situations, improving the physiological states of the friends and enhancing their survival [14–16]; and friendships, even same-sex, can positively influence reproductive outputs [17, 18]. Although the evolutionary implications are well established, the proximate mechanisms of animal friendships are relatively understudied. Consequently, empirical evidence on the behavioural decision rules that apply in a ‘biological market’ to choose preferred partners or friends is lacking [19].

Several behavioural decision rules have been proposed as proximate mechanisms of animal friendships [20], of which the ‘homophily principle’ received much attention. The homophily principle, proposed originally to characterise human social networks, suggests that individuals similar in terms of their age, gender, ethnicity, and interests are more likely to become friends [21]. To some extent, these propositions fundamentally overlapped with that of another behavioural decision rule, the symmetry-based reciprocity [22, 23]. Yet, friendships in animals can form irrespective of age, sex, and kin relationships, and thus, these variables could not solely explain the emergence of friendships [24]. Personality – the consistent inter-individual differences [25] – due to their immense adaptive values [26], have been proposed as an alternate mechanism influencing friendships. Similarities or differences in traits are calculated at the dyadic levels to evaluate how personalities may influence friendships. Interestingly, when extended to personalities, researchers have found strong evidence of homophily explaining the emergence and sustenance of friendships in both humans [27, 28] and animals (chimpanzee [29], bonobo [30], Assamese macaque [31], baboon [32]). Given that these findings are restricted to (non-human-)primates, testing the hypothesis in phylogenetically diverse taxa is required to ascertain if personality homophily is a general behavioural decision rule of friendship.

Ungulates represent a wide range of species with varying social organisations [33], providing a suitable system to expand the investigation of the emergence of friendships. Several ungulate species exhibit complex sociality patterns, including marked preferences for specific group members over others (e.g., cattle [34], goat [35], Przewalski’s horse [36], bison [37]). However, existing research has predominantly used demographic characteristics, e.g., age and sex, often coupled with kinship and dominance rank relationships, to explain sociality patterns in ‘non-random’ or heterogeneous networks. In recent years, studies have assessed the personality traits of various ungulate species and identified their influence on cognitive performance and decision-making [38, 39], autonomic nervous system reactivity [40], dominance hierarchy [41], and welfare [42], among others. To our knowledge, no systematic attempt has been made to investigate the effects of personality homophily on the emergence of ungulate heterogeneous social networks, including friendships. Moreover, most research on ungulate social behaviour and personality is performed in captivity, often as livestock, thus partly limiting the broader ecological value of the findings [43].

Water buffalo (synonymously, buffalo) are domesticated ungulates with an estimated global population of 208 million [44]. Almost all research on buffalo is restricted to their production capacities (for meat and milk) as livestock. In feral and free-living conditions, female buffalo can form clans of up to 30 individuals, comprised of kin members (grandmothers, mothers, daughters, and sisters), whereas multiple such non-kin or distantly related clans can form herds of up to 500 individuals; Males typically disperse and form bachelor groups of up to 10 individuals [45]. However, our understanding of their social interaction patterns is very limited. Despite this considerable gap in knowledge, some research, predominantly conducted in captivity, suggests that buffalo are highly social animals, exhibiting complex patterns of sociality [46], sociability and affiliative behaviours [46, 47], and dominance-rank relationships [48]. Thus, to truly understand their social behaviour, it is necessary to carry out research on populations that are largely free of human interventions, living in wild or feral states. Such natural populations can also prove to be valuable in investigating the effects of personality homophily on social associations.

Here, we studied a feral and free-ranging population of buffalo. To assess friendships in female buffalo, we utilised two behavioural indicators: proximity and affiliative body contact [20]. Unlike non-human primates and many ungulate species, allogrooming, a strong indicator of social bonds, is predominantly absent in buffalo [49]. In species with limited direct affiliative interactions (e.g., allogrooming), spatial associations (e.g., proximity and body contact) can sufficiently capture the varying social bonds among individuals [50]. We conducted extensive behavioural observations and objectively assessed the consistent inter-individual differences or personalities in female buffalo. Notably, we used a ‘bottom-up’ approach to assess personalities; hence, traits were not predetermined [51–54]. We hypothesised that if friendships among female buffalo are formed based on personality homophily, then homophily in personality is a general behavioural decision rule for partner preference. In particular, we predicted lower score differences in social personality traits would be positively associated with higher dyadic proximity values.

## Results

### Friendships in buffalo

We validated the dyadic proximity index, i.e., our measure of buffalo social associations, by comparing it with the dyadic body contact index (see **Methods**). We found a moderately strong positive correlation (Spearman rank correlation: n=274, ρ = 0.42, p < 0.001), suggesting that the dyadic proximity index sufficiently captured affiliative social relationships (**Figure 1**). On average, the proximity index had a value of one (range = 0.15 to 2.19), with higher values indicating stronger social associations. For instance, a dyadic value of 2.19 indicates 2.19 times stronger associations between two individuals than the population average.

**Figure 1.**
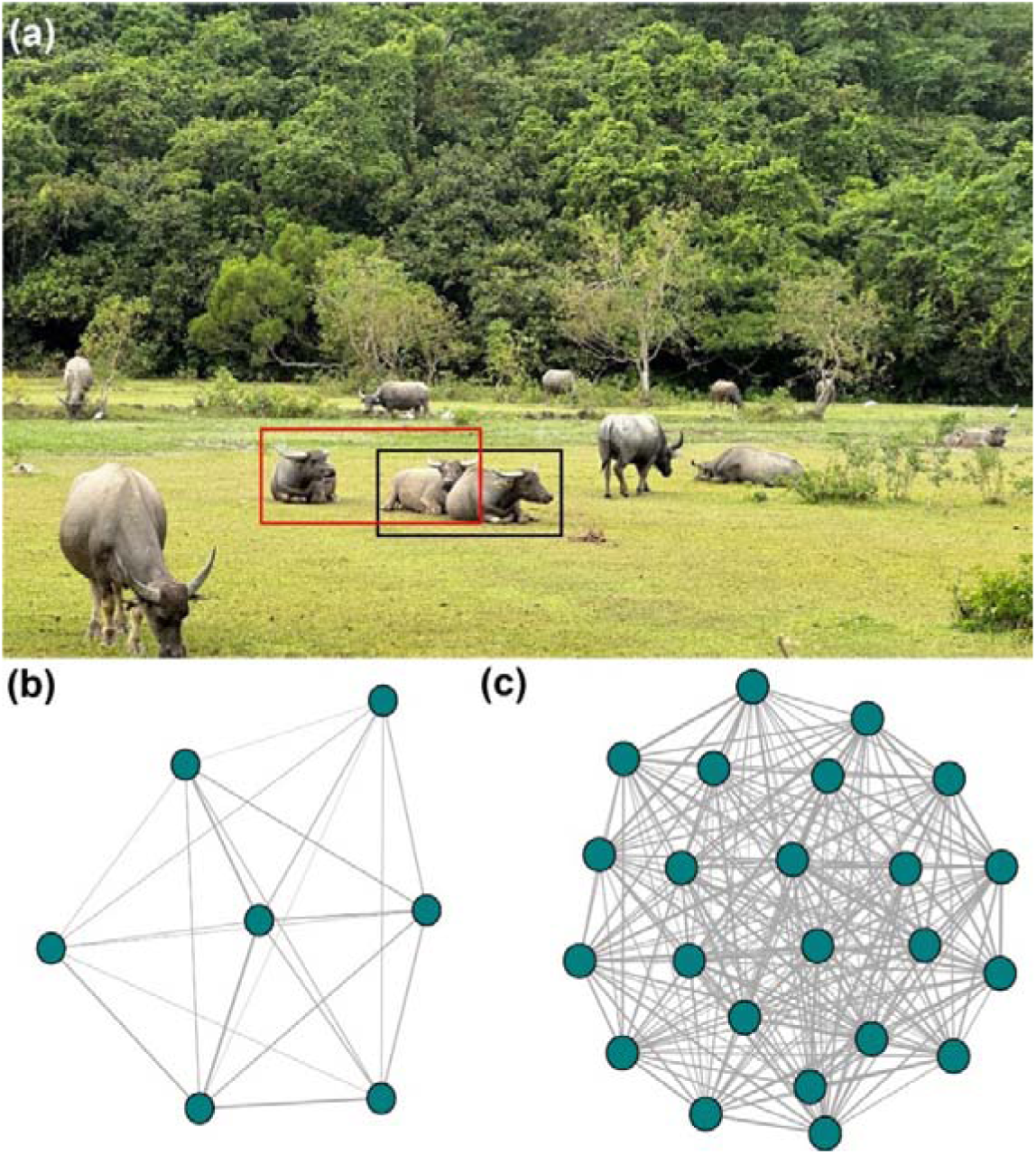
The spatial positions of females in a feral and free-ranging population of buffalo. **(a)** The black rectangle shows two female buffalo sitting in affiliative body contact. The red rectangle shows two female buffalo sitting in proximity (one body length distance); **(b)** Proximity network of female buffalo in the Lo Wai Tsuen herd; **(c)** Proximity network of female buffalo in the Lo Uk Tsuen herd. In (b) and (c), each dark green circle or node indicates an individual, and the lines or edges between them indicate the dyadic proximity index values, with thicker edges indicating higher index values or strong social associations. [Photo credit (a): Debottam Bhattacharjee, Location: Pui O, Hong Kong].

### Personality traits in buffalo

We extracted three principal components (or personality traits) from a principal component analysis (PCA) that included six repeatable behavioural variables (**Figure 2**, also see **Methods**). The three PCs cumulatively explained 80.15% of the variance (**Table S1**). The three PCs were labelled based on the behavioural variables that were loaded on them (**Table S2**). PC1 had three positively loaded variables: *approach*, *self-groom*, and *avoid*. We labelled it as *social tension*. PC2 included only one variable, *vigilance*; hence, it was given the same label, i.e., *vigilance*. PC3 had two loaded variables, *sit* and *displace*, loaded positively and negatively, respectively. We labelled PC3 *general dominance*.

**Figure 2.**
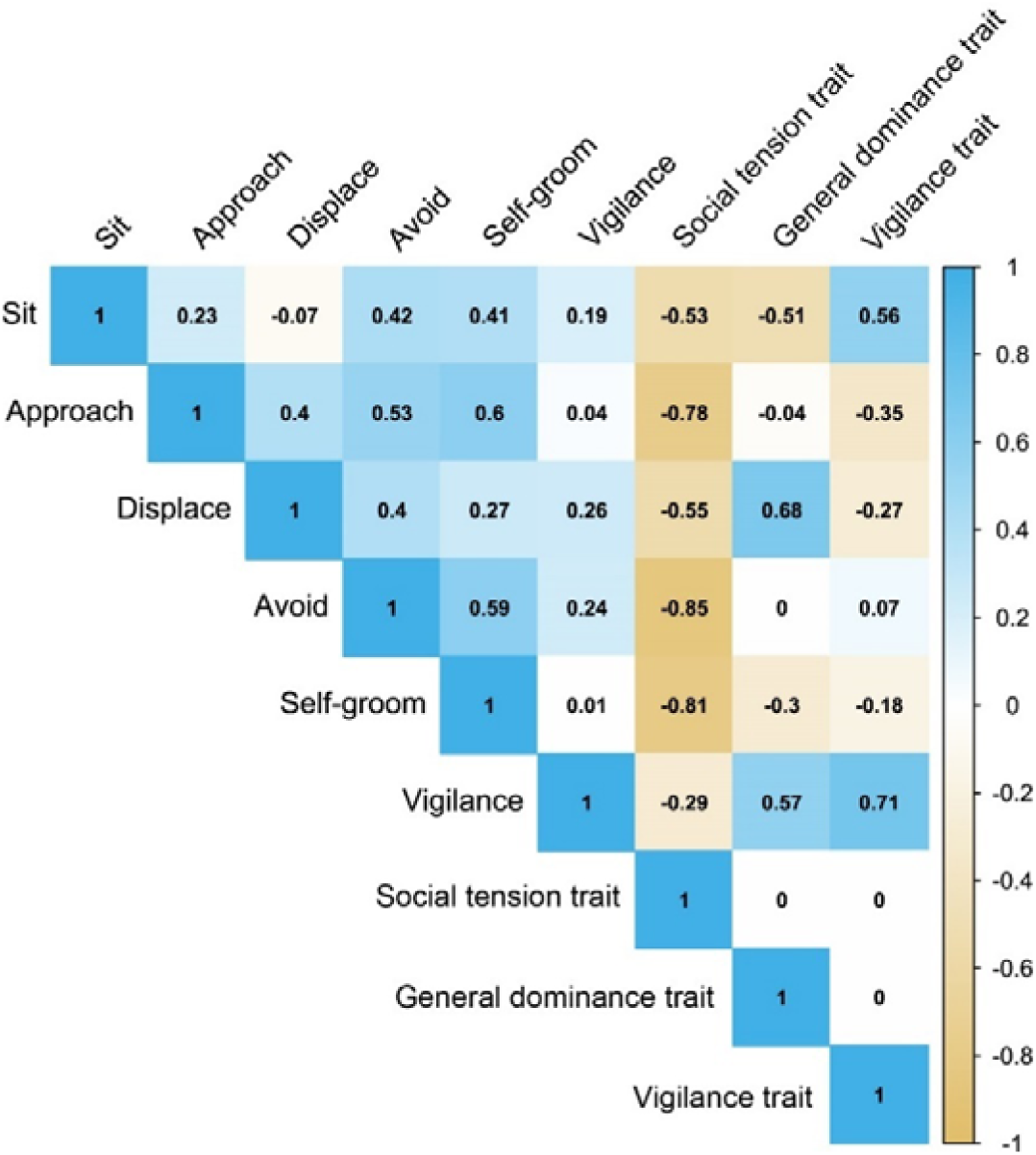
Correlation matrix plot of behavioural variables and buffalo personality traits. From each PC or personality trait, individual factor scores (synonymously personality scores) were extracted, and score differences were calculated for all combinations of dyads. As personality traits are often considered age-independent behavioural constructs, we tested for correlations between individual age and personality scores for the three traits separately. Personality traits and age had no significant relationships (Spearman rank correlation tests, social tension: ρ = -0.01, p = 0.93, vigilance: ρ = -0.01, p = 0.92, general dominance: ρ = 0.006, p = 0.97).

### Effects of personality traits, age differences, kinship, and rank differences on buffalo friendships

We found significant relationships between personality trait differences and the dyadic proximity index (**Figure 3**, **Table S3**). Lower dyadic score differences in social tension were associated with higher dyadic proximity index values (GLMM: z = -5.147, Cohen’s d = 0.62, 95% CI = [0.39, 0.86], p <0.001, **Figure 3a**). Differences in vigilance did not affect the dyadic proximity index (GLMM: z = 0.768, p = 0.44, **Figure 3b**). Like social tension, dyads with lower score differences in general dominance had higher proximity index values (GLMM: z = -4.427, Cohen’s d = 0.53, 95% CI = [0.30, 0.77], p <0.001, **Figure 3c**). We did not find any effect of age differences on the dyadic proximity index (GLMM: z = 1.468, p = 0.14). Similarly, kinship had no effect on the dyadic proximity index (GLMM: z = 1.110, p = 0.26, **Figure 3d**).

**Figure 3.**
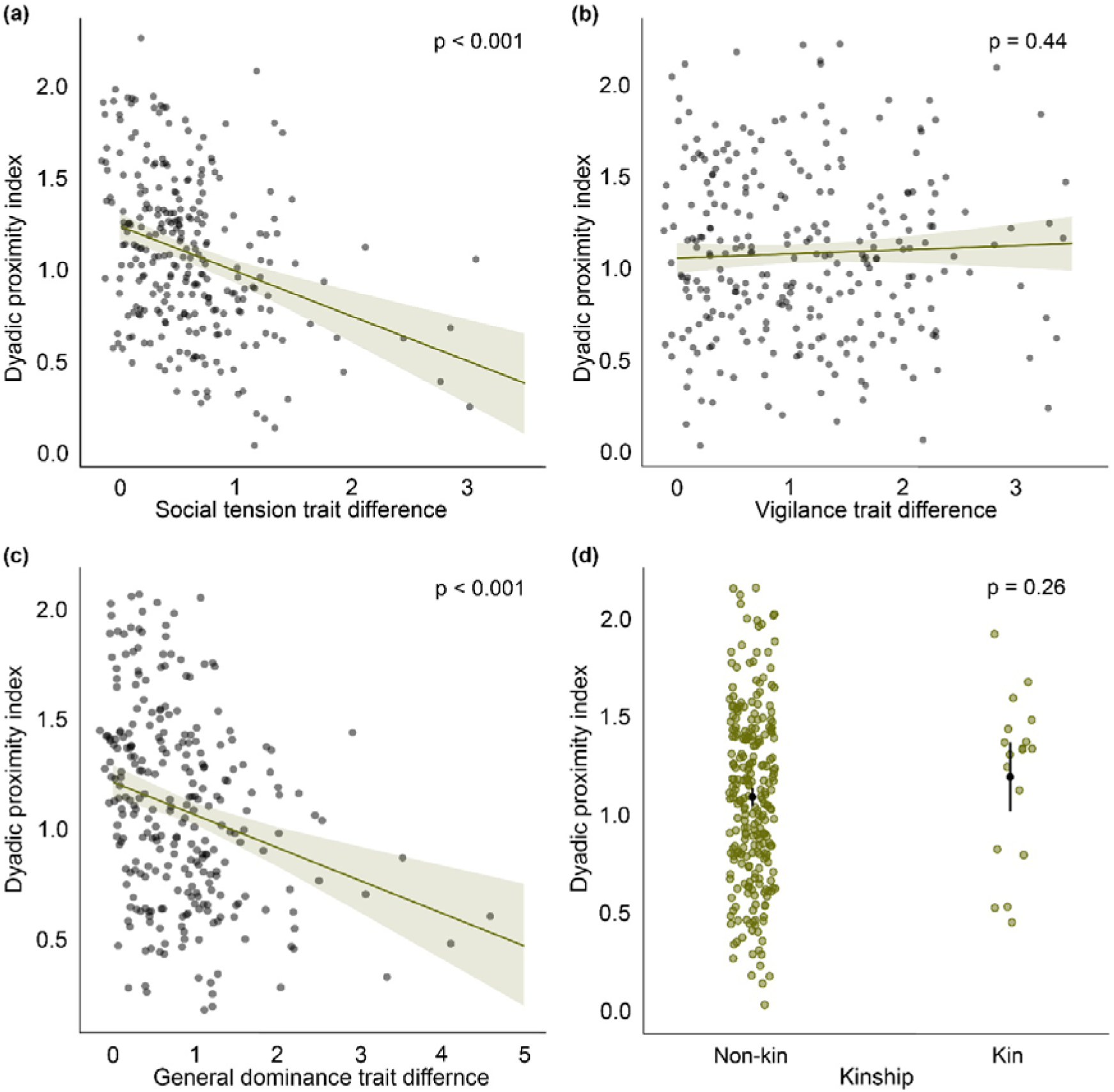
Predicted effects of buffalo personality trait differences and kinship on dyadic proximity index. **(a)** Effect of social tension trait differences on proximity index; **(b)** Effect of vigilance trait differences on proximity index; **(c)** Effect of general dominance trait differences on proximity index; **(d)** Effect of kinship on proximity index. Solid dark green circles indicate data points. In (a), (b), and (c), shaded areas indicate 95% confidence intervals. In (d), solid black dots and vertical lines indicate predicted mean values and 95% confidence intervals, respectively.

These results suggest that dyads with more similar social tension and general dominance traits were more likely to be in close proximity than dyads with greater trait differences. This full model differed from the null model (Likelihood ratio test: ^2^ = 57.918, p <0.001). We found no collinearity among the fixed effects (VIF range: 1.01 - 1.12), and the model residuals followed normality assumptions.

Some behavioural variables (i.e., avoid and displace), which constitute social tension and general dominance personality traits, are often used in combination with other behaviours, such as agonistic interactions, to construct dominance hierarchies [55]. However, we found that these direct interactions were very rare during scans, leading to > 94% ‘unknown’ dominance rank relationships among the individuals. Thus, dominance hierarchies could not be constructed based on the data collected during scans. Yet, to investigate whether dominance hierarchies were associated with friendships, we assessed hierarchies based on our overall continuous focal data. The buffalo herds had low to moderately steep dominance hierarchies (Steepness: Lo Wai Tsuen = 0.61 ± 0.07, Lo Uk Tsuen = 0.53 ± 0.04). However, we did not find any effect of dominance rank differences on dyadic proximity index (GLMM: z = -0.166, p = 0.87, **Table S4**).

## Discussion

The emergence of friendships in animals has long been considered a phenomenon governed by similarities in demographic characteristics, such as age and sex [4]. However, the sustenance of age- and sex-independent friendships has impelled researchers to propose alternate explanatory mechanisms, like homophily in personality [29]. Yet, empirical research explicitly testing homophily as a behavioural decision rule of friendship is mainly restricted to primates. We investigated adult same-sex friendships in a free-ranging and feral population of a social ungulate species, the water buffalo [56]. Our results show that non-random female-female social associations form in feral buffalo, where stronger associations may indicate friendships [20], and homophily in certain personality traits can predict them. Similarities in age and dominance rank relationships did not explain the dyadic social associations. Also, we found that kin and non-kin dyads show comparable levels of dyadic associations. Based on these results, we show that homophily in personality is a proximate behavioural decision rule of friendship that applies to species beyond primates.

### Friendships in buffalo

The dyadic proximity index, validated by affiliative body contact, captured the differential strengths of buffalo social associations, including friendships. Friendships in buffalo can have considerable socio-ecological and adaptive implications, which can range from foraging and collective movement to resting. The ‘conspecific attraction hypothesis’ suggests that the presence of conspecifics can be used as social cues during foraging site selection, especially when resources are patchily distributed [57] (also see [58]). However, social aggregations during foraging are often considered a by-product of sharing space (i.e., random associations) and not outcomes of preferential associations. Although empirical testing of such assumptions is limited, conscious behavioural synchronisation [59] or exhibition of the same behaviour by strongly bonded individuals in different species is evident. Synchronisation of activities, like foraging and collective movement, in a multi-level society of feral horses is influenced by dyadic social associations, with higher associations leading to better synchronisation [60]. Stronger affinitive bonds result in more successful spatiotemporal movements in horses, too [61]. Desert baboons follow their friends during the departure phase of collective movements [62]. In addition, microhabitat-level space use patterns, in terms of resting site selection, can be governed by preferential strong social associations. Cape buffalo and forest buffalo choose specific resting sites (e.g., forest clearing) that enable them to sit in very close proximity to each other (often including body contact) and facilitate social interactions [63–65]. Close spatial associations can foster allopreening in parrots and corvids [66]. These results strongly suggest that preferential social associations can form during foraging, collective movements and resting, and such associations may not necessarily be random and simple by-products of sharing a common space. Therefore, while the general herd-level social associations in space use patterns may strengthen group cohesion and help avoid predators through ‘many eyes’ [67], preferential social associations or friendships can still be sustained. As evident in a species like water buffalo, where direct affiliative interactions, like allogrooming, are predominantly absent [49], spatial proximity patterns can be of vital importance for maintaining friendships (cf. [50]). Hence, instead of fully relying on the assumption of the “gambit of the group” [68], species-specific ‘norms’ of friendships can be present, i.e., without necessitating direct affiliative interactions among friends (cf. [20, 29, 49, 50, 69]).

Our assessment of buffalo friendships encompassed all aspects of grazing, movement, and resting in dry and wet seasons. The strength of overall observed differential dyadic associations suggests that friendships were likely sustained throughout contexts and climatic conditions. Friendships were both age- and kin-independent. Kinship can govern close social associations potentially mediated through ‘familiarity’. Nonetheless, friendships may form both within genetically closely related and unrelated members, suggesting that kinship is not necessarily a prerequisite for the emergence of friendships [12, 20, 29]. Additionally, dominance rank differences did not predict friendships in buffalo. In highly despotic societies with steep dominance hierarchies, rank-based cooperation among group members or selectivity in benefiting group members is evident [12, 70]. The low to moderate steepness values in dominance hierarchies in female buffalo might explain why rank differences are not suitable drivers of friendships.

### Personality traits in buffalo

Using an objective bottom-up method, we reported three personality traits in feral buffalo: social tension, vigilance, and general dominance. While these personality traits are subjectively labelled (cf. [30, 41, 51–54]), the underlying repeatable behaviours may have socio-ecological implications for buffalo. The social tension trait consisted of three behaviours: *approach*, *self-groom*, and *avoid*. Approaching a conspecific or even heterospecific and tolerating them is interpreted as the tendency to be sociable [71], which has substantial adaptive implications (see [43] for a detailed discussion). Self-groom, or broadly, self-directed behaviour in animals, is a widely used indicator of stress-related responses, contributed by social (e.g., hierarchical group structure) or non-social (e.g., presence of ectoparasites) factors [52, 72–74]. Finally, avoiding (specific) conspecifics can be attributed to the hierarchical social structure of buffalo (see [48]), which may help reduce conflicts, competition and disease transmission within the herd [75]. However, counterintuitively, approach behaviour positively co-varied with self-groom and avoidance (of conspecifics) in our personality construct, indicating aspects of both sociability and tension. The availability of partners, space, and distribution of food resources in social and ecological niche can potentially explain why behaviours of two extremes may have positively co-varied [76]. In other words, individuals may show consistent behavioural ‘tactics’ to fulfil their social and ecological needs.

The second buffalo personality trait, vigilance, was labelled after the only behavioural variable loaded in it. Vigilance, in general, has a high adaptive value associated with predator avoidance; however, our study population of feral buffalo has no natural predators. Thus, proximately, this behaviour can be attributed to focusing on within-herd events, like ongoing physical fights, or paying attention to nearest neighbours [77]. In addition to resident males within the herds, other territorial males who were not part of the observed herds were present in the study area. Subsequently, frequent territorial fights among resident and non-resident males, often followed by female guarding, were observed (personal observations, D.B., 2023-2024). The approach of the resident male for mate guarding can make nearby females vigilant. However, individuals who are not within a guarding radius can avoid being vigilant. Further, individuals with relatively low dominance ranks can collect spatiotemporal information on aversive events (like fights and aggression) and nearest neighbours (like an approaching male or other higher-ranking herd members) and benefit by escaping or avoiding conflicts. Therefore, consistent variations in vigilance behaviour in feral buffalo can provide substantial benefits even in the absence of natural predators.

The third and final personality trait of buffalo, general dominance, has two inversely co-varying behaviours: *sit* and *displace*. In the context of water buffalo, the behaviour *sit* can extend beyond resting. The swamp and marshland habitats allowed our study population to ‘rest’ in the waterlogged fields. While the behaviour was coded independently of wallowing [78], sitting in such a terrain may include attributes of thermoregulation, ectoparasite removal, etc, thus potentially serving valuable physiological functions. However, such semi-naturally occurring waterlogged areas are not uniformly distributed [79], especially during the dry season of the year, leading to competition (including *displace* behaviours) among individuals for access. Therefore, the inversely co-varying behaviours can have consistent inter-individual differences and be labelled general dominance. Besides, this trait is often linked to boldness and exploration [80] (but see [81]), which are highly beneficial when predator pressure is low or absent. In contrast, individuals with low general dominance scores can benefit by avoiding potential conflicts and aggressive interactions. General dominance has further been reported as a manifestation of leadership behaviour in animals [82], which could be helpful in decision-making, e.g., in collective movements. To what extent variations in general dominance represented the dominance hierarchy of the herds remains to be investigated, which could not be assessed in this study to avoid data dependence.

### Personality homophily and friendships

As hypothesised, we found evidence that personality homophily is a behavioural decision rule of friendship in female water buffalo. Social tension and general dominance, but not vigilance, predicted the preferential strong social associations. Unlike vigilance, social tension and general dominance are social personality traits (i.e., traits that include social interactions [83]). In general, social personality traits, like extraversion and agreeableness, are known to foster friendships and cooperation in humans [84]. Certainly, the buffalo social personality traits do not fully resemble the human social personality traits, yet justification can be made by highlighting the underlying basis of personality homophily: its ability to help form trust and reduce uncertainty among similar individuals [4, 21]. Consistent with this idea, similarly (socially-) tensed buffalo can form emotionally mediated attitudes toward herd members through social associations, leading to preferential selection of partners [52, 85]. Likewise, differences in general dominance have significant implications for adaptive benefits. A low dominance rank difference or high dominance rank similarity can be associated with better cooperation and coordination [12, 86]. Friendships based on similar general dominance status can reduce the monopolisation of resources like food, by facilitating tolerance. In line with previous findings, the non-social vigilance trait did not influence friendships [31], which may suggest that the emergence of friendships in buffalo may depend on social but not on non-social personality traits (but see [29]).

## Conclusion

The evolution of social relationships has several complex underpinnings beyond simple demographic characteristics. Using an underrepresented social ungulate species, we provide valuable insights into the proximate mechanisms driving preferential close social relationships or friendships. We identify and discuss the emergence of friendships based on similarities in personalities, a behavioural decision rule which has so far been tested primarily on primates. However, the involved psychological and emotional processes employed for the selection of friends in water buffalo should carefully be assessed before generalising them with that of humans and other non-human primates. Although we highlight the broader evolutionary implications of friendships in buffalo by drawing parallels with other social species, significant gaps in knowledge of feral water buffalo ecology and behaviour have rendered identifying the species-specific norms of sociality and friendships challenging. These limitations call for in-depth investigations to validate how the broad spectrum of ecological relevance pertaining to a host of other taxa aligns specifically with water buffalo.

## Methods

### A. Study area and population

We conducted this study on a feral and free-ranging population of water buffalo present in Pui O in the Southern Lantau Island of Hong Kong SAR, China. Pui O is a small village at the edge of Lantau South Country Park, and the marshlands of Lo Wai Tsuen (22°14’32.7" N 113°58’42.3" E) and Lo Uk Tsuen (22°14’29.3" N 113°58’31.1" E) in Pui O are home to two different herds of feral buffalo. At the beginning of the study, the Lo Wai Tsuen herd had a size of 19 (females = 7, males = 12), whereas the Lo Uk Tsuen herd comprised 29 individuals (females = 25, males = 4), of which two females were below two years of age (i.e., calves). We collected data on thirty adult females (age range: 4 to 19; mean age in years ± standard deviation = 10.38 ± 4.33) from the two different herds (Lo Wai Tsuen = 7, Lo Uk Tsuen = 23, **Table S5**) between 17^th^ July 2023 and 6^th^ February 2024, including a wet and a dry season [87]. While the number of females remained the same throughout the study period, the number of males changed in both herds due to death and dispersal. Two male buffalo from the Lo Wai Tsuen herd dispersed in October, and one individual died in late December (due to natural causes). One male buffalo from the Lo Uk Tsuen herd dispersed in January. Although the two herds live adjacently, inter-herd interactions among females were never observed (personal observations, D.B., 2023-2024, also see [88]). The buffalo fed on grass and other natural vegetation but occasionally (three to four times a week between December and February) received supplementary food (hay and sweet potato leaves) from local citizen groups. Fresh water was available *ad libitum* from small waterbodies in and around the marshlands.

A cattle management team from the Agriculture, Fisheries and Conservation Department of Hong Kong (AFCD) routinely sterilised and managed the buffalo population [89]. All adult females were sterilised except one from the Lo Uk Tsuen herd, and their status did not change during the study period. While over 50% of the adult females (16 of 30) had numbered ear tags (administered by the AFCD), we relied on morphological features, such as horn shape and structure, relative horn length and scar marks, to identify the remaining individuals. A photo catalogue was prepared to ensure the reliable identification of the individuals. Kinship and individual age details were collected from a local non-government citizen group, documenting the buffalo population’s demographic information over the past 20 years.

### B. Ethical declarations

Ethical approval was obtained from the Animal Research Ethics Sub-Committee of [Anonymous]. The behavioural observations of buffalo were conducted from a distance of at least 20 m without direct human intervention. We adhered to the ethical guidelines of the ASAB/ABS for conducting the research [90].

### C. Data collection

Continuous focal and scan sampling methods were used to collect behavioural data [91]. Each observation day was divided into three time periods: morning (0900-1159 hours), afternoon (1200-1459 hours), and late afternoon (1500-1800 hours), and each period was equally sampled. An extensive ethogram was developed with all behaviours exhibited by the buffalo (**Table S2**), which was used during data collection and coding.

#### Focal sampling

Each continuous focal observation session was 20 minutes long. We collected 720.03 ± 0.32 minutes (mean ± standard deviation) of focal data per individual. Thus, each individual was observed a total of 36 times. We used a semi-randomised order to observe the individuals in different time periods on an observation day, and the same individual was not observed more than once per day. We conducted one to nine focal observations daily (5.17 ± 1.96 focal observations/day), at least three times a week. Focal data were recorded using a video camera (Panasonic HC-V785) mounted on a tripod or collected live using digital data sheets. All state (durations) and point (frequencies) behaviours of the focal individuals, as well as their interactions with non-focal herd members, were noted using the ethogram.

#### Herd scan sampling

We followed predetermined routes during scan sampling. The sampling distances of Lo Wai Tsuen and Lo Uk Tsuen were 2.1 km and 2.3 km, respectively. The same routes had to be walked more than once to cover the sampling area, but data were recorded only for the first time. During each scan, we recorded the spatial positions of the individuals relative to each other in a herd. We collected data on the frequency of two levels of dyadic spatial positions: dyadic proximity (one body length distance to each other) and affiliative body contact (body parts of two individuals excluding legs or horns touching while standing, sitting, or lying down) (**Figure 1a**, **Table S2**). We conducted 3 – 5 scans per day (3.84 ± 0.74 scans/day) at least twice a week. The interscan interval was 30 minutes, and we conducted 300 scans of each herd. Notably, all individuals from their respective herds were present during all the scans.

### D. Data analyses and statistics

#### Coding

Videos were played using Pot Player (version 240315, 1.7.22129) and coded in a frame-by-frame manner. Following the ethogram, we coded the duration (in seconds) and frequency of all relevant state and event behaviours, respectively (cf. **Table S2**). Since each individual was observed for the exact duration (i.e., 720 minutes), no time correction was deemed necessary. D.B. coded all the videos and compiled them with the focal data recorded in digital datasheets. Another person, unaware of the goals of the study, coded 5% and 10% of the focal and scan data, respectively. The reliability between D.B. and the blind coder was excellent (Intraclass correlation tests: focal data - ICC (3,k) = 0.93, p < 0.001; scan data - ICC (3,k) = 0.95, p < 0.001).

#### Assessment of friendships

We created all possible combinations of dyads at the herd level (Lo Wai Tsuen herd = 21 dyads and Lo Uk Tsuen herd = 253 dyads). Four of the 21 dyads from the Lo Wai Tsuen herd consisted of kin members, whereas only 14 were kin dyads out of the 253 dyads from the Lo Uk Tsuen herd. Kinship specifically included mother-daughter, grandmother-granddaughter, and sibling (i.e., sisters) relationships. From the herd scans, the frequencies of proximity and affiliative body contact were used to calculate a dynamic dyadic sociality index (DSI) [92, 93], a widely used indicator of strong social associations and friendship in animals. Dyadic proximity and body contact frequencies were divided by the respective population mean values to calculate corresponding index (i.e., dyadic proximity index and body contact index) values. These scaled index values were combined and averaged to obtain the DSI values. However, the data on the frequency of body contact was zero-inflated, with only close to 30% of the potential dyads exhibiting affiliative body contact. To avoid any statistical model convergence issues, we chose dyadic proximity index as the only indicator of friendship (see [20, 69]). Nevertheless, we investigated the validity of dyadic proximity index by comparing it with the body contact index (**Figure 1a**).

#### Assessment of personality

A standardised ‘bottom-up’ approach was used to assess the personality traits [51–53], where any potential bias of predetermined clustering of variables could be avoided. By definition, personality traits should be consistent across time and contexts [25]; therefore, we divided the focal data into two phases (with 360 min/focal individual for each phase) to investigate whether they were repeatable. The repeatability of behavioural variables was assessed using a two-way mixed model intraclass correlation (ICC (3,1)) test [51–53]. Forty-eight behavioural variables (cf. **Table S2**) were considered for the bottom-up approach. However, variables with low frequencies were dropped before conducting the ICC analysis.

Twenty-seven variables were dropped as more than half of the individuals did not exhibit them, i.e., the variables had zero occurrences. Subsequently, twenty-one variables were retained for the ICC analysis (**Table S6**). We used a conservative ICC cut-off value of 0.5 for a variable to be considered repeatable [94]. Fourteen of the twenty-one variables were repeatable, with ICC values ranging from 0.521 to 0.850 (**Table S6**).

A principal component analysis (PCA) was conducted using the fourteen repeatable variables. The average values of the repeatable variables between the two phases were calculated, standardised, and included in the PCA. However, due to low Kaiser-Meyer-Olkin sampling adequacy (MSA) values (MSA cut-off value = 0.6) and inadequate percentage communality values (communality cut-off value = 70%) [95], the number of variables was further reduced to six by a step-by-step process (cf. [51, 52]). The percentage of communalities of the remaining six variables ranged between 73.02 to 91.33, with an overall MSA value of 0.69. A scree plot was generated using an unrotated PCA, and the eigenvalue of each potential principal component (PC) was retrieved. The number of PCs was subsequently decided based on the eigenvalues (with eigenvalue ≥1) and by visual inspection of the scree plot (**Figure S1**) [95]. All other assumptions of PCA were met (Bartlett test: p < 0.001, and cumulative variance explained by PCs > 60%). As personality traits can be correlated and form behavioural syndromes [96], we used an oblique rotation technique (direct oblimin rotation). The factor loadings ≥ 0.6 (positive and negative) were considered significant [97].

#### Effects of personality trait differences on friendships

A generalised linear mixed-effect model (GLMM) analysis was conducted to investigate the effects of personality score- and age-differences and kinship on dyadic proximity index values. We used a Gaussian error distribution with an ‘identity’ link function. In the full model, the dyadic score differences of each personality trait, absolute dyadic age differences, and kinship (yes/no) were included as fixed effects and the proximity index as a response variable. Individual identities of a dyad (i.e., individual 1 and individual 2) were included as random effects. The null model included the response variable and the random effects but lacked fixed effects.

#### Assessment of dominance hierarchies and their effects on friendships

Using focal data, we created four frequency-based matrices per herd where all individuals were placed as initiators and receivers of avoid, displace, physical aggression, and flee behaviours (see **Table S2** for definitions). Subsequently, we built combined matrices, each for a herd, that followed consistent directional patterns of the behaviours (i.e., avoid, being displaced, being physically aggressed, and flee). We used a Bayesian Elo-rating method to assess herd-level hierarchies [98]. The method involved the calculation of winning probabilities from the combined matrices.

Upon construction, individuals were plotted ordinally along their respective hierarchies based on the estimated Elo values. The ordinal rank differences were calculated for the dyads and standardised to account for the varying herd sizes. Since personality and dominance data are not independent, to avoid any bias of data dependence, rank difference was not included as a fixed effect in the GLMM that investigated the effects of age and personality differences on friendships. In a separate GLMM with a Gaussian error distribution and ‘identity’ link function, dyadic rank difference was included as a fixed effect and the proximity index as a response variable. Individual identities of a dyad were included as random effects. The null model included the response variable and the random effects but lacked the fixed effect.

#### Statistical packages

All statistical analyses were performed in R (version 4.3.1) [99]. ICC analyses were conducted using the *psych* package [100]. We used *Elosteepness* package to construct the dominance hierarchies [98]. We used the package *glmmTMB* for the GLMM analyses [101]. The null vs full model comparisons were checked with the *lmtest* package [102]. Collinearity among fixed effects was investigated using the *performance* package [103], and a variation inflation factor (VIF) of >3 was considered a threshold for high correlation [104]. GLMM model diagnostics (normality assumptions) were checked using the package *DHARMa* [105]. The significance value was set at 0.05 for all statistical tests.

## Acknowledgements

We thank Wong Ching Ki and Tin Ka Yuen for their help with data collection and coding. We sincerely thank Jean Leung for providing the demographic data on the buffalo. We are grateful to Jorg Massen, Elham Nourani, George Hodgson, and Tania Perroux for their valuable feedback on an earlier version of this manuscript.

## Authors’ contributions

D.B.: conceptualisation, methodology, investigation, data curation, formal analysis, data visualisation, writing—original draft, writing—review and editing; K.J.F.: conceptualisation, funding acquisition, resources, supervision, writing—review and editing; A.G.M.: conceptualisation, funding acquisition, resources, supervision, writing—review and editing. All authors gave final approval for publication and agreed to be held accountable for the work performed therein.

## Data availability

Data and code will be made available upon publication.

