## Supplementary Material for "Personality homophily drives female friendships in a feral ungulate"

*Electronic supplementary materials to*

**Table S1. The rotated principal components or buffalo personality traits with variable loadings and their percentages of variance, and percentages of attribute communalities of the behavioural variables.**

| **Variables** | **Social tension** | **Vigilance** | **General dominance** | **% Communalities** |
| --- | --- | --- | --- | --- |
| Approach | 0.87 |  |  | 73.12 |
| Self-groom | 0.86 |  |  | 77.27 |
| Avoid | 0.76 |  |  | 73.02 |
| Vigilance |  | 0.97 |  | 91.33 |
| Sit |  |  | 0.80 | 85.98 |
| Displace |  |  | -0.64 | 83.62 |
| **% Variance** | 41.30 | 20.10 | 18.75 |  |

Variable loadings of >±0.6 are considered significant.

**Table S2 – Ethogram describing the buffalo behaviours and their descriptions.** The behavioural categories, names of behaviours, their types and definitions are provided.

| **Category** | **Behaviour** | **Type** | **Definition** |
| --- | --- | --- | --- |
| Activity | Walk | State | Individual moves at a regular pace, without ruminating, foraging or grazing and covers at least a three-body length distance. |
|  | Stand | State | Individual stands upright without moving, ruminating, or paying attention to a particular object, event or another individual. |
|  | Sit | State | Individual sits with her head up but without ruminating. |
|  | Lie down | State | Individual lies with one side of the body on the ground. The head position is always down. |
|  | Ruminate | State | Individual chews stored food items with lateral jaw movements, head above mid-line or in line with the body while standing, sitting, or lying. |
|  | Feed | State | Individual places her mouth at the level of grass with clear visible contact. Movement of the mouth can be visible. |
|  | Browse | State | Individual stands with her mouth visibly in touch with plant leaves or shrubs but not grass. Movement of the mouth can be visible. |
|  | Supplement feed | State | Individual feeds on human-provisioned food which are otherwise not part of the natural vegetation, e.g. sweet potato leaves and/or hay. |
|  | Drink | State | Individual places her mouth at the level of water with clearer movements indicative of swallowing water. |
|  | Sleep | State | Individual lies with head down and eyes closed. |
|  | Pass by | Event | Individual enters the radius of one body length of another individual and leaves the radius with the same movement. |
|  | Defecate | Event | Individual releases faeces, usually in a squatting position. |
|  | Urinate | Event | Individual releases urine, often in a semi-squatting position. |
|  | Startle | Event | Individual shows frightened response (frozen body posture with attention towards stimuli) due to a sudden change in the environment (e.g., loud sound, other members moving or loud noises with human artefacts). |
|  | Digging wallow | State | Individual digs soil with her horns and/or hooves with the intention of making a depression to accommodate herself in mud and/or water. |
|  | Wallow | State | Individual is present within a wallow, which can happen in conjunction with sitting, lying down, and ruminating. |
|  | Bath | State | Individual is present within a natural waterbody, can happen in conjunction with standing, sitting, lying, and ruminating; the individual can be fully submerged. |
| Sexual behaviour | Mount | Event | Individual lifts her front side of the body and places it against another individual (usually the hind side of the individual) with two feet off the ground. |
|  | Flehmen | Event | Individual exposes her nostrils and teeth to inhale olfactory cues from another individual; This may pair with lifting the neck and extending the chin with a slightly open mouth and upper lip curled back backwards. |
|  | (Being) Mate | State | Individual mounts and ejaculates, indicated by a pause before dismounting and/or the appearance of visible semen. (For females: Mate is ‘being mate’ by a male, i.e., a passive behaviour) |
|  | Refusal | Event | Individual leaves or changes position, with or without aggression, while another individual tries to mount. |
| Social behaviour | Approach | Event | Individual moves toward another individual with clear focus and ends up within their one-body length radius. |
|  | Social rub | State | Individual rubs her body gently on a conspecific’s body without displacing the conspecific. |
|  | Social sniff | Event | Individual sucks in the air while holding her head very close to body parts of another individual, except for the genital area. |
|  | Lick | Event | Individual passes tongue or lips over the body of another individual, except the genital area. |
|  | Head touch with head/muzzle | Event | Individual touching the head of another individual with her own head/muzzle. |
|  | Touch other body parts than head with head/muzzle | Event | Individual touching body part except for head of another individual with own head/ muzzle. |
|  | Contact sit or stand (body contact) | State | Individual sits or stands next to another individual with body parts other than horns and limbs touching. |
|  | Contact lie down (body contact) | State | Individual lies next to another individual with body parts other than horns and limbs touching. |
|  | Proximity | State | Individual stays within one body length of another individual. |
|  | Social play | Event | (Either of) Two individuals skip/hop with each other and nuzzle, which do not end in withdrawal. |
|  | Muzzle contact | Event | Individual brings her mouth close to another individual’s mouth region. |
|  | Vocalisation | Event | Individual emits vocal sound directed at another individual/human/event with head directed towards the point of interest; can also happen during walking. |
|  | Travel together | State | Individual travels parallel to another individual while being within a three body length radius. |
|  | Follow | Event | Individual walks behind another individual in the same direction and stays within a radius of five body lengths. Behavioural synchrony is present. |
| Agonistic interactions | Displace | Event | Individual causes non-contact movement of another individual away from their previous position, taking at least three steps. |
|  | Avoid | Event | Individual changes her sitting spot or direction while walking to a distance of at least three body lengths from another individual that approaches the focal. Attention of the focal is on the approaching individual. |
|  | Leave | Event | Individual walks away from another individual who is within a radius of one body length. |
|  | Flee | Event | Individual runs away after being approached/chased/lunged at by another individual. |
|  | Freeze | Event | Individual stops displaying body movements (i.e., ear or tail movements) and stands stiff while another individual passes by. Duration is a maximum of 20 seconds(considered another event when exceeds 20 seconds); can happen while being inspected by another individual. |
|  | Physical fight | State | Individuals engages in consistent physical contact with another individual acting to displace the other; the outcome is separation of the two individuals, often with a clear winner. |
|  | Push | Event | Individual forces to change the position of another individual by contact, often with horns. |
|  | Head butt | Event | Individual swings her head with force against the body of another individual. |
| Self-directed behaviour | Autogroom | Event | Individual licks self with tongue in contact with body, or self-scratches using limbs or inanimate objects such as trees or fence; includes parts such as head, ears, neck, shoulders, back, flanks, rear, legs, belly, and others. |
|  | Head/body shake | Event | Individual rapidly turns the whole body in at least two different directions. |
|  | Yawn | Event | Individual opens her mouth and inhales intensely, which can be seen by the expansion of the chest. |
|  | Vigilance | Event | Individual focuses on an event or another individual without body movements for at least three seconds, often exposing nostrils and teeth for inhaling olfactory cues (if focusing on an individual). |
| Interspecific interactions |  | States and events | Individual directly interacts with a member of a different species (humans, cattle, dogs). Due to the rarity of occurrences, all possible behaviours are pooled. |

**Table S3 – Generalised linear mixed-effect model analysis investigating the effects of buffalo personality- and age- differences and kinship on dyadic proximity index.**

**Model:** Dyadic proximity index ~ Social tension difference + Vigilance difference + General dominance difference + Age difference + Kinship + (1|Individual 1) + (1|Individual 2), family=Gaussian.

|  | **Estimate** | **Standard error** | **Z value** | **p-value** |
| --- | --- | --- | --- | --- |
| Intercept | 1.291 | 0.058 | 22.251 | <0.001*** |
| Social tension difference | -0.244 | 0.047 | -5.147 | <0.001*** |
| Vigilance difference | 0.021 | 0.027 | 0.768 | 0.442 |
| General dominance difference | -0.149 | 0.033 | -4.427 | <0.001*** |
| Age difference | 0.009 | 0.006 | 1.468 | 0.142 |
| Kinship (yes) | 0.102 | 0.092 | 1.110 | 0.267 |

Significance code: p <0.001 ‘***’

**Table S4 – Generalised linear mixed-effect model analysis investigating the effects of buffalo dominance rank differences on dyadic proximity index.**

**Model:** Dyadic proximity index ~ Dominance rank difference + (1|Individual 1) + (1|Individual 2), family=Gaussian.

|  | **Estimate** | **Standard error** | **Z value** | **p-value** |
| --- | --- | --- | --- | --- |
| Intercept | 1.083 | 0.034 | 31.20 | <0.001*** |
| Dominance rank difference | -0.004 | 0.027 | -0.166 | 0.868 |

Significance code: p <0.001 ‘***’

**Table S5 – Details of the study animals.** The individual identities, herd memberships, age in years at the beginning of the study (June 2023), and dominance ranks are summarised.

| **Identity** | **Herd membership** | **Age in years** | **Dominance rank** |
| --- | --- | --- | --- |
| Ross’s Mum | Lo Wai Tsuen | 19 | 2 |
| Siu Ying Mum | Lo Wai Tsuen | 19 | 1 |
| Siu Ying | Lo Wai Tsuen | 6.5 | 3 |
| Siu Ying Sis 1168 | Lo Wai Tsuen | 6 | 4 |
| Chu 802 | Lo Wai Tsuen | 10 | 5 |
| Chu Daughter | Lo Wai Tsuen | 6 | 7 |
| Weak Sister Ping tag | Lo Wai Tsuen | 6 | 6 |
| 857 | Lo Uk Tsuen | 15 | 23 |
| 856 | Lo Uk Tsuen | 14 | 20 |
| 855 | Lo Uk Tsuen | 14 | 18 |
| Ear Pink tag | Lo Uk Tsuen | 15 | 19 |
| Chase Car | Lo Uk Tsuen | 12 | 17 |
| Chase Car Daughter | Lo Uk Tsuen | 5 | 14 |
| Amy's Mum | Lo Uk Tsuen | 14 | 8 |
| Cloud | Lo Uk Tsuen | 6 | 13 |
| 1160 | Lo Uk Tsuen | 6 | 22 |
| Baby Girl's mum 812 | Lo Uk Tsuen | 10 | 21 |
| Baby Girl 1097 | Lo Uk Tsuen | 5 | 6 |
| Older Sister 1076 | Lo Uk Tsuen | 9 | 1 |
| Younger Sister | Lo Uk Tsuen | 9 | 3 |
| Yellow Button | Lo Uk Tsuen | 14 | 11 |
| Sneaky Boy's mum | Lo Uk Tsuen | 15 | 16 |
| Fighter Girl 1166 | Lo Uk Tsuen | 4 | 10 |
| Ah Mui Yellow Button | Lo Uk Tsuen | 6 | 4 |
| 1158 | Lo Uk Tsuen | 8 | 5 |
| 1045 | Lo Uk Tsuen | 9 | 7 |
| 1159 | Lo Uk Tsuen | 15 | 2 |
| No Hair Pink Tag | Lo Uk Tsuen | 14 | 12 |
| 1035 | Lo Uk Tsuen | 11 | 9 |
| 1091 | Lo Uk Tsuen | 9 | 15 |

**Table S6. Behavioural variables from the buffalo ethogram used in the intraclass correlation analysis and their repeatability measures.** Intraclass correlation (ICC (3,1)) analysis output of behavioural variables with their ICC values, 95% confidence intervals (CI) and F- and p-values.

| **Variables** | **ICC** | **95% CI** | **F-value** | **p-value** |
| --- | --- | --- | --- | --- |
| Walk | 0.170 | -0.197, 0.495 | 1.410 | 0.180 |
| Stand | 0.241 | -0.125,0.549 | 1.635 | 0.090 |
| Sit | **0.693** | 0.448, 0.841 | 5.511 | <0.001 |
| Lie down | **0.580** | 0.284, 0.776 | 3.765 | <0.001 |
| Ruminate | -0.146 | -0.477, 0.220 | 0.745 | 0.784 |
| Feed | 0.499 | 0.174, 0.725 | 2.989 | 0.002 |
| Pass by | 0.402 | 0.055, 0.662 | 2.344 | 0.013 |
| Wallow | **0.521** | 0.204, 0.739 | 3.176 | 0.001 |
| Bath | 0.312 | -0.048, 0.601 | 1.907 | 0.044 |
| Flehmen | **0.722** | 0.494, 0.857 | 6.202 | <0.001 |
| Approach | **0.710** | 0.474, 0.850 | 5.886 | <0.001 |
| Contact sit | **0.619** | 0.338, 0.799 | 4.250 | <0.001 |
| Contact lie down | 0.479 | 0.150, 0.713 | 2.840 | 0.003 |
| Proximity | **0.579** | 0.282, 0.775 | 3.754 | <0.001 |
| Travel together | **0.554** | 0.247, 0.759 | 3.482 | 0.001 |
| Displace | **0.601** | 0.312, 0.788 | 4.009 | <0.001 |
| Avoid | **0.728** | 0.503, 0.861 | 6.355 | <0.001 |
| Leave | **0.592** | 0.300, 0.783 | 3.906 | <0.001 |
| Self-groom | **0.850** | 0.709, 0.926 | 12.32 | <0.001 |
| Head/Body shake | **0.788** | 0.602, 0.893 | 8.446 | <0.001 |
| Vigilance | **0.777** | 0.583, 0.887 | 7.974 | <0.001 |

Variables with ICC values ≥0.5 (highlighted by bold interface) are considered repeatable.

**Figure S1 –**

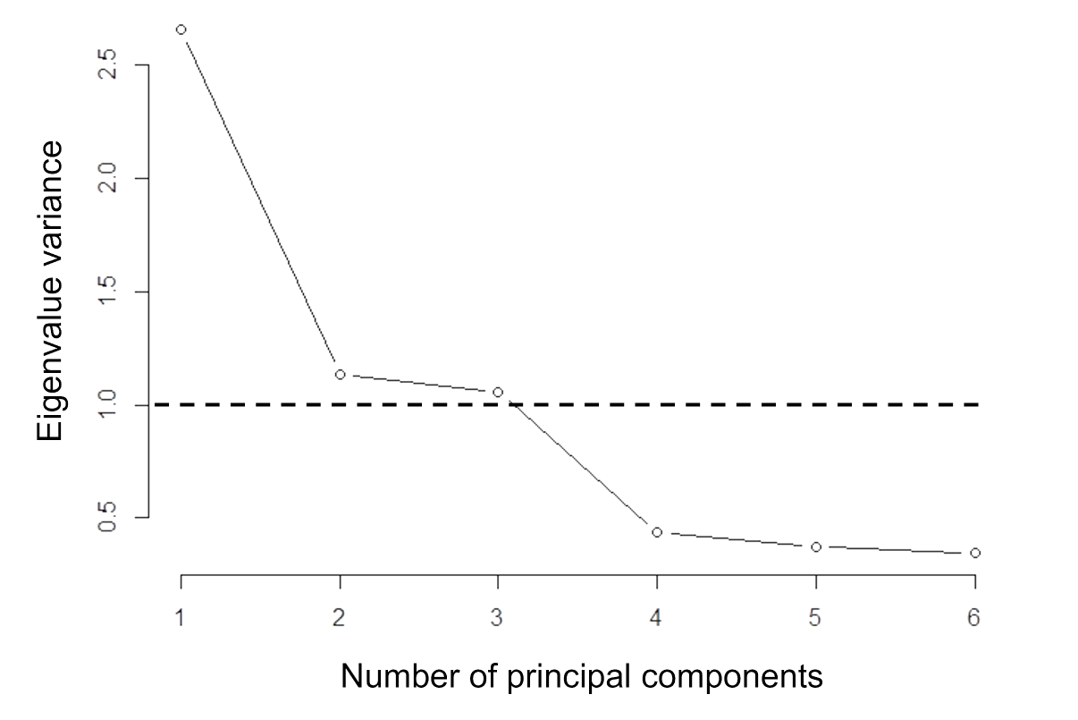

**Figure S1 – Screeplot of the principal component analysis to identify buffalo personality traits.** The horizontal dashed line indicates the cut-off eigenvalue of ‘1’ for selecting a principal component. Three principal components are selected with eigenvalues >1.
